# Activation of the hypothalamic-pituitary adrenal axis in response to a verbal fluency task and associations with task performance

**DOI:** 10.1101/2019.12.30.890814

**Authors:** Linda Becker, Ursula Schade, Nicolas Rohleder

## Abstract

Speech fluency can be impaired in stressful situations. The aim of the present study was to investigate whether a verbal fluency task without any further stress induction by itself induces responses of the hypothalamic-pituitary adrenal (HPA) axis and of the sympathetic nervous system (SNS).

The sample consisted of *n* = 85 participants (68.2% female; 33.3 ± 15.2 years, BMI = 23.7 ± 4.3 kg/m^2^) who performed two consecutive verbal fluency tasks for two minutes each. The categories were either ‘stress’ or ‘disease’ and ‘animals’ or ‘foods’ which were presented in a randomized order. Three saliva samples were collected, prior to the task (t_0_), immediately after (t_1_), and ten minutes after (t_2_). Salivary α-amylase and cortisol were assessed. Furthermore, blood pressure, heart rate, and subjective ratings of actual stress perception, level of effort, and tiredness were measured.

The verbal fluency task induced an HPA axis response with a maximum cortisol level at t_2_ (*p* < .001, η_p_^2^ = .19) which was independent of task performance. Furthermore, subjective stress and effort as well of tiredness increased immediately after the task (all *p* < .001; all *d* ≥ 1.0). Tiredness immediately after the task was negatively correlated with task performance (*p* = .002). No α-amylase, blood pressure or heart rate responses were found.

We conclude that a verbal fluency task acts like an acute stressor that induces a cortisol response without the need of further (e.g., social-evaluative) stress components. Therefore, it is a little time-consuming alternative to other stress tasks that can be used in field studies with little effort.

## Introduction

Acute stress triggers to a variety of physiological responses. The most prominent is the activation of the hypothalamus-pituitary adrenal (HPA) axis which leads to secretion of the stress hormone cortisol from the adrenal cortex. Furthermore, the sympathetic nervous system (SNS) is activated in response to acute stress which leads to the release of epinephrine and norepinephrine from the adrenal medulla, as well as to a variety of secondary reactions such as an increase in blood pressure, heart rate, and a decrease in heart rate variability (1–3). Both stress systems can interact with the brain via direct and indirect pathways and can, therefore, alter brain chemistry which can consecutively alter cognitive functioning (4–6). For example, retrieval from declarative long-term memory decreases after an acute stressor and this is associated with the HPA axis response (7–9). Furthermore, working memory is affected in response to an acute stressor which is associated with both the HPA axis and the SNS response (10–12). A cognitive function that has been less investigated so far, is verbal fluency (VF) which describes the ability to name as many terms that belong to a specific rule (e.g., from a specific category or words that begin with a specific letter) as possible during a predefined time interval. Verbal fluency tasks (VFT) require a variety of cognitive processes such as long-term memory and working memory as well as executive functions (13,14). It has been found previously that speech productivity can be impaired in acute stress situations. Buchanan and colleagues (2014, 15) found that participants with high cortisol responses to the Trier Social Stress-Test (TSST; 16), paused more during the stress task than participants with low cortisol responses. However, it is still an open question whether a speech task itself can induce a stress response.

Therefore, the aim of the present study was to investigate whether performing a VFT without any further stress induction, by itself, leads to a physiological stress response. If one was found, we were further interested in whether the stress response was associated with cognitive performance during the task. Furthermore, we aimed to investigate – if a response was found – whether this is associated with subjective stress perception and mood, as well as with anthropometric and health factors (e.g., age, sex, BMI, and depression) that are typically related with HPA axis and SNS responses.

## Materials and Methods

### Participants

From initially *n* = 101 (67/66.3% female; 34.9 ± 15.2 years; BMI = 23.7 ± 4.0 kg/m^2^; 81/80.2% non-smokers) participants, ten (9.9%) were excluded because they did not provide enough saliva for analysis, five (5%) because German was not their mother tongue, and one (1%) because his performance in the VFT was below three standard deviations from the sample’s mean. The final sample consisted of *n* = 85 healthy participants (58/68.2% female; 33.3 ± 15.2 years, min.: 18, max.: 69; BMI = 23.7 ± 4.3 kg/m^2^, min.: 18.2, max.: 41.5; 71/83.5% non-smokers). About half of the participants (48/56.5%) came to our laboratory during a public event in the evening on a weekend between 6 p.m. and 1 a.m. The other sessions took part on weekdays between 9 a.m. and 5 p.m. For the morning sessions, the participants were instructed to have gotten-up at least two hours before the start of the experiment. Participants refrained from drinking (except water) and eating within at least one hour prior to the experiment. All participants gave their written and informed consent. The study was conducted according to the principles expressed in the Declaration of Helsinki and was approved by the local ethics committee of the Friedrich-Alexander University Erlangen-Nürnberg (# 397_19 B).

### General procedure

Participants were seated in a comfortable chair in a quiet room. After they were informed about the experimental procedure, they gave their written consent for participation, and the first saliva sample (s_0_) was collected, and blood pressure and heart rate were measured for the first time. The VFT (see below) was then introduced to the participants and started immediately. After this, the second saliva sample was collected and both blood pressure and heart rate were measured. To fill the gap between the second and third saliva collection, participants filled-out questionnaires, assessing their demographic variables, health status, chronic stress perception, and lifestyle factors (see below for a more detailed description). Ten minutes after s_1_, the last saliva sample (s_2_) was taken and blood pressure and heart rate were measured again. If necessary, participants were given more time to complete the questionnaires after s_2_. Finally, weight and height were measured. During collection of the saliva samples, participants rated their actual stress level, level of effort, and tiredness (see below). The time course of the whole experiment is shown in Fig 1. The whole session lasted between 20 and 30 minutes, depending on the time needed for informing the participants and for filling-out the questionnaires.

**Fig 1.**
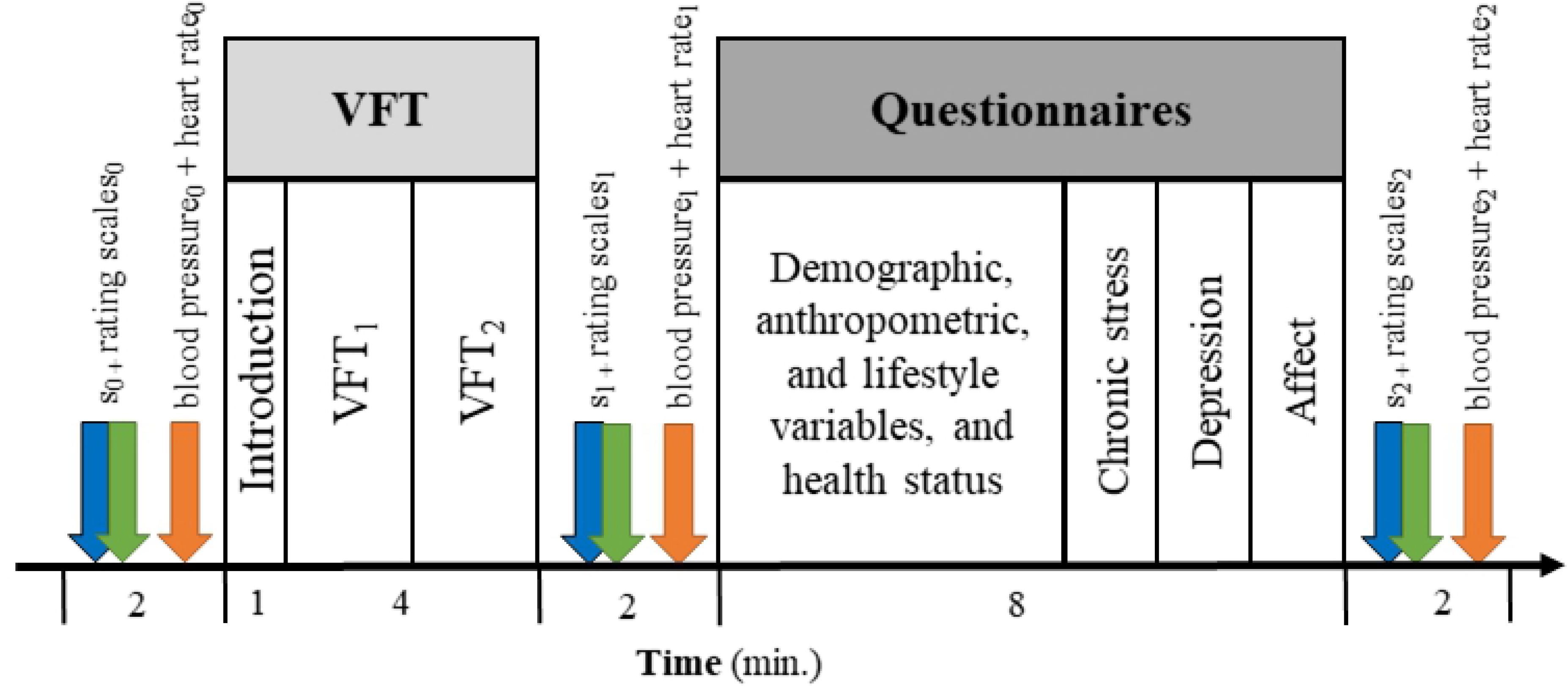
Time course of the experiment. Three saliva samples s_0_, s_1_, and s_2_ were collected. The verbal fluency task (VFT) was performed between the first and the second saliva sample and consisted of two parts (VFT_1_ and VFT_2_). Furthermore, blood pressure was measured after each saliva sample.

### Speech fluency assessment

A category fluency task was used for assessing speech fluency. The participants were given a category for which they should name as many terms as possible. Each participant was assigned two categories, one neutral and one emotional, and they were given two minutes time for each. The neutral category was either “animals” or “foods” and the emotional category was either “stress” or “disease”. The order of the categories was counterbalanced between the participants. Participants were voice recorded during the VFT, using a digital dictaphone (Olympus VN-541PC). For VFT evaluation, the number of correct terms (VFT_corr,i_), number of repetitions (VFT_rep,i_), and number of other errors (VFT_oth,i_, e.g. wrong category) were determined and a performance score VFT_perf,i_ = VFT_corr,i_ – (VFT_rep,i_ + VFT_oth,i_) was calculated for each category i = {1, 2}. From these, a mean performance score VFT_perf_ = (VFT_perf,1_ + VFT_perf,2_)/2 was calculated.

### Physiological assessment

#### Saliva sampling and analysis

Saliva samples were collected by means of salivettes (Sarstedt, Nümbrecht, Germany). Participants were instructed to keep the salivette in their mouth for at least one minute and to move it back and forth, but not to chew on it. Saliva samples were stored at −30 °C after collection for later analyses. Immediately before analysis, samples were centrifuged at 2000 g and 20 °C for ten minutes. Salivary α-amylase (sAA) was measured with an in-house enzyme kinetic assay using reagents from DiaSys Diagnostic Systems GmbH (Holzheim, Germany), as previously described (17,18). In brief, saliva was diluted at 1:625 with ultrapure water, and diluted saliva was incubated with substrate reagent (α-amylase CC FS; DiaSys Diagnostic Systems) at 37° C for three minutes before a first absorbance reading was taken at 405 nm with a Tecan Infinite 200 PRO reader (Tecan, Crailsheim, Germany). A second reading was taken after five minutes incubation at 37 °C and increase in absorbance was transformed to sAA concentration (U/ml), using a standard curve prepared using “Calibrator f.a.s.” solution (Roche Diagnostics). Salivary cortisol concentrations were determined in duplicate using chemiluminescence immunoassay (CLIA, IBL, Hamburg, Germany). Intra- and inter-assay coefficients of variation were below 10% for both sAA and cortisol.

#### Blood pressure and heart rate

Systolic and diastolic blood pressure as well as heart rate were assessed by means of an upper arm blood-pressure monitor (boso medicus X). Two participants (2.4%) were excluded from blood pressure and heart rate analysis because of technical problems. Twenty-five participants (29.4%) were classified as hypertonic because their blood pressure during the t_0_ measurement was ≥140/90 mmHg. From these, only two (8%) reported a hypertension diagnosis. The hypertonic participants were initially excluded from analysis of the blood pressure and heart rate time course, but further analyses were performed that confirmed that the same effects were found for hypertonic and non-hypertonic participants. Since blood pressure might be related to the other physiological variables as well and, therefore, with the sAA or cortisol response, we investigated whether the same sAA and cortisol time courses were found for hypertonic and non-hypertonic participants. However, no differences were found as well (see below).

### Further assessment

#### Subjective ratings

During saliva collection, subjective stress, level of effort, and tiredness were rated on 10-point Likert scales by the participants. The anchors were “not stressed at all” and “extremely stressed”, “no effort” and “extreme effort”, as well as “not tired” and “extremely tired”.

#### Demographic variables, health status, and lifestyle factors

Self-developed questionnaires were used to assess demographic variables (age, sex, BMI, marital status, educational level, occupation, monthly income), health status (physiological and psychological diseases, medication) and lifestyle factors (smoking, alcohol consumption, participation in regular sports). Body-mass index was classified according to the norms provided by the World Health Organization (WHO) as underweight (< 18.5 kg/m^2^), normal weight (18.5 – 24.9 kg/m^2^), pre-obese (25 – 29.9 kg/m^2^), and obese (>29.9 kg/m^2^).

#### Chronic stress

The perceived level of chronic stress within the last four weeks was assessed by means of a German translation of the 10-item version of the Perceived Stress Scale (PSS; 19).

#### Depression

Depression was assessed by means of the German version of the long form of the depression scale from the Center for Epidemiological Studies (CES-D; 20,21). A cut-off value of 22 was used for classification into depressed and non-depressed participants (22).

#### Mood

As last part of the questionnaire battery and immediately before the last saliva sample, actual mood was assessed by means of a German version of the positive and negative affect schedule (PANAS; 23–25).

### Statistical data analysis

For statistical analyses, IBM SPSS Statistics (version 26) was used. Normality of distribution was tested by means of the Kolmogorov-Smirnov test. Because of positive skewness and violation of normality, sAA levels were transformed by means of the square root transformation and cortisol levels by means of the natural logarithm prior to further statistical analysis. Analyses of variance for repeated measurements (rmANOVAs) with the within-subject factor time (t_0_, t_1_, t_2_) were calculated, separately for subjective stress ratings, sAA, and cortisol levels as well as for blood pressure and heart rate. For subjective stress ratings, the within-subject factor ‘state’ with the levels ‘stress’, ‘effort’, and ‘tiredness’ was included. For blood pressure a within subject factor ‘component’ with the levels ‘systole’ and ‘diastole’ was included. To correct for multiple comparisons and because six separate rmANOVAs were calculated, an adjusted α-level of α_adjusted_ = α/6 = 0.05/6 = 0.008 was used. Partial eta-squares (η_p_^2^) were considered as effect sizes. Sphericity was tested by means of the Mauchly test (26). If necessary, degrees of freedom were corrected by means of the Greenhouse-Geisser procedure (27). For post-hoc analysis, *t*-tests for dependent samples were calculated and Cohen’s *d* was considered as measure for effect sizes. For these dependent *t*-tests, Cohen’s *d* was corrected according to the method that was proposed by Morris (2008, 28). One-factorial ANOVAs were used to investigate whether performance differences exist between the different VFT orders or in the physiological and subjective stress variables between the BMI groups. For exploratory analyses, Pearson correlations between age, BMI, depression, chronic stress, and time of day and the physiological and subjective stress variables were calculated. An adjusted α-level of α_adjusted_ = 0.05/27 = 0.002 was used because 27 correlations were calculated (cortisol, sAA, stress, effort, tiredness, heart rate, systole, diastole at three time points as well as positive and negative affect, and VFT_perf_). Furthermore, it was examined whether differences in group means between depressed and non-depressed, high and low chronically stressed, and between hypertonic and non-hypertonic participants can be found. For this, independent t-tests with the same adjusted α-level of α_adjusted_ = 0.002 were calculated.

Since response to the VFT were found for cortisol between t_0_ and t_2_ and for the subjective ratings between t_0_ and t_1_, percentage increases were calculated for these variables as follows: cort_increase_ = (cort_t2_ – cort_t0_)*100/cort_t0_ and rating_increase_ = (rating_t1_ – rating_t0_)*100/rating_t0_. The data set that was used for statistical analysis and the corresponding codebook can be found in the Supplementary Files S1 and S2.

### Power analysis

An a-priori power analysis was performed, using G*Power (version 3.1.9.2). It indicated an optimal sample size of *n*_optimal_ = 82. Power analysis was performed for an rmANOVA with an α-level of α = .05, a power of 1 - β = .95, an effect size of *f* = 0.2, three groups (e.g., for subjective ratings), and three measurement time points (t_0_, t_1_, and t_2_). Therefore, we assume that the achieved sample size of *n* = 85 is sufficient for the reported analyses.

## Results

### Sample description

Most (62/72.9%) of the participants reported that they regularly engaged in exercise on 3 ± 1.5 days per week on average. Most (35/56.5%) were doing endurance sports, three (4.8 %) strength training, three (4.8%) relaxation training, and 21/33.9% a combination of endurance, strength, and relaxation training. According to the cut-off value by Stein and Luppa (2012; 22), ten participants (11.8%) were classified as depressive. Thirty-eight (44.7%) participants were classified as low and 47/55.3% as high chronically stressed. An overview of the other relevant descriptive sample characteristics is provided in Table 1.

**Table 1.** Sample characteristics.

| Variable | Label | Number | Percent age |
| --- | --- | --- | --- |
| Sex | Female | 58 | 68.2 |
|  | Male | 27 | 31.8 |
| BMI classification | Underweight | 2 | 2.4 |
|  | Normal weight | 61 | 71.8 |
|  | Pre-obese | 15 | 17.6 |
|  | Obese | 7 | 8.2 |
| Marital Status | Single | 39 | 45.9 |
|  | Married | 23 | 27.1 |
|  | Partnership | 21 | 24.7 |
|  | Divorced | 1 | 1.2 |
|  | Widowed | 1 | 1.2 |
| Education | Secondary school | 4 | 4.7 |
|  | Medium maturitary | 11 | 12.9 |
|  | Professional High School | 5 | 5.9 |
|  | A-levels | 35 | 41.2 |
|  | Bachelor | 13 | 15.3 |
|  | Master, diploma | 11 | 12.9 |
|  | PhD | 5 | 5.9 |
|  | Habilitation | 1 | 1.2 |
| Profession | Unemployed | 2 | 2.4 |
|  | Trainee | 5 | 5.9 |
|  | Student | 31 | 36.5 |
|  | Employed | 36 | 42.4 |
|  | Self-employed | 1 | 1.2 |
|  | Official | 6 | 7.1 |
|  | Pension | 2 | 2.4 |
|  | Other | 2 | 2.4 |
| Smoking | Yes, daily | 7 | 8.2 |
|  | Yes, sometimes | 7 | 8.2 |
|  | No | 71 | 83.5 |
| Alcohol consumption on study day | Yes, but less than 2 beverages and not within the last 2 hours | 11 | 12.9 |
|  | No | 74 | 87.1 |
| General alcohol consumption | No | 10 | 11.8 |
|  | 1 day | 16 | 18.8 |
|  | 2 days | 11 | 12.9 |
|  | 3 days | 6 | 7.1 |
|  | 4 days | 1 | 1.2 |
|  | 5 and more days | 3 | 3.5 |
|  | Less than 1 day per week | 38 | 44.7 |
| Use of oral contraceptives (women only) | No | 47 | 81 |
|  | Yes | 11 | 19 |

### Verbal fluency performance

The mean VFT performance was VFT_perf_ = 26.9 ± 7.3 (min.: 10.5, max: 48.5). Mean VFT performances for each category are summarized in Table 2. Mean performances did not differ between the different presentation orders and category combinations (stress/animals, stress/foods, disease/animals, disease/foods, animals/stress, animals/disease, foods/stress, and foods/disease; *p* = .925). Verbal fluency performance did not differ between the stress and disease category (*p* = .063) and between the animals and foods category (*p* = .230).

**Table 2.**
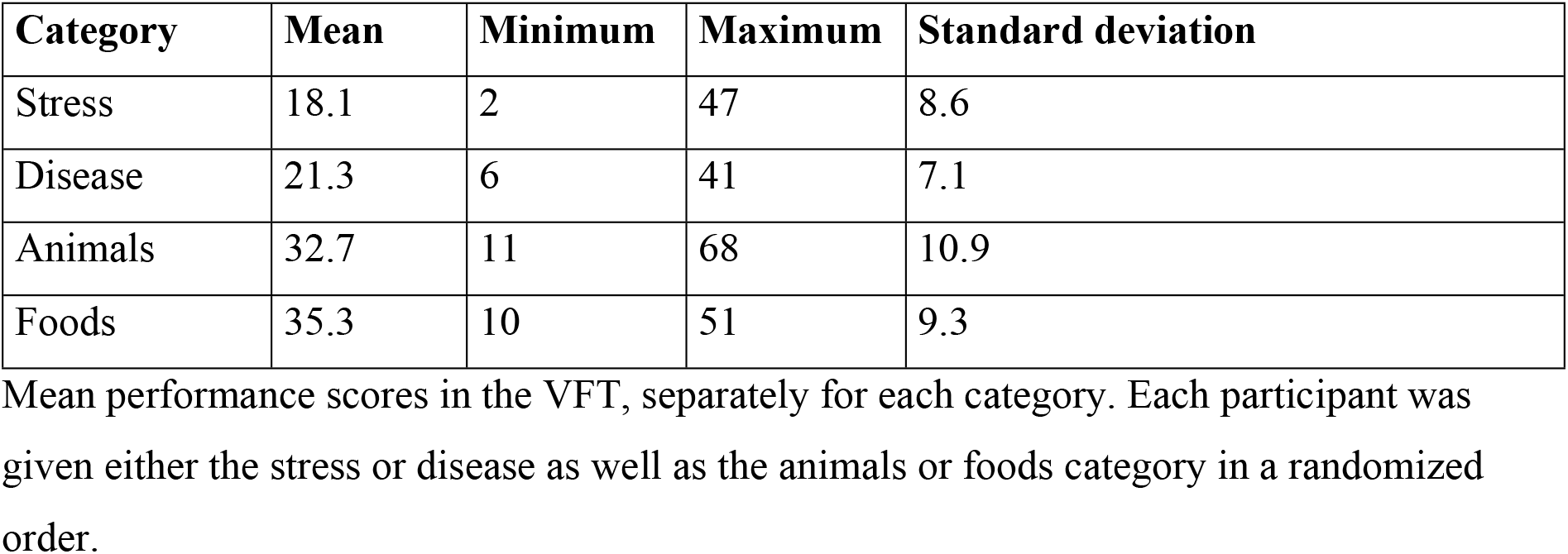
Verbal fluency task performance.

| Category | Mean | Minimum | Maximum | Standard deviation |
| --- | --- | --- | --- | --- |
| Stress | 18.1 | 2 | 47 | 8.6 |
| Disease | 21.3 | 6 | 41 | 7.1 |
| Animals | 32.7 | 11 | 68 | 10.9 |
| Foods | 35.3 | 10 | 51 | 9.3 |

Mean performance scores in the VFT, separately for each category. Each participant was given either the stress or disease as well as the animals or foods category in a randomized order.

### Alpha-amylase

Salivary α-amylase levels did not significantly differ between the three measurement-time points (*p* = .487, η_p_^2^ = .008; Fig 2a) and were not associated with VFT performance (all *p* ≥ .293). When only including the non-hypertonic participants into the analysis, also no effects were found (all *p* ≥ .057).

**Fig 2.**
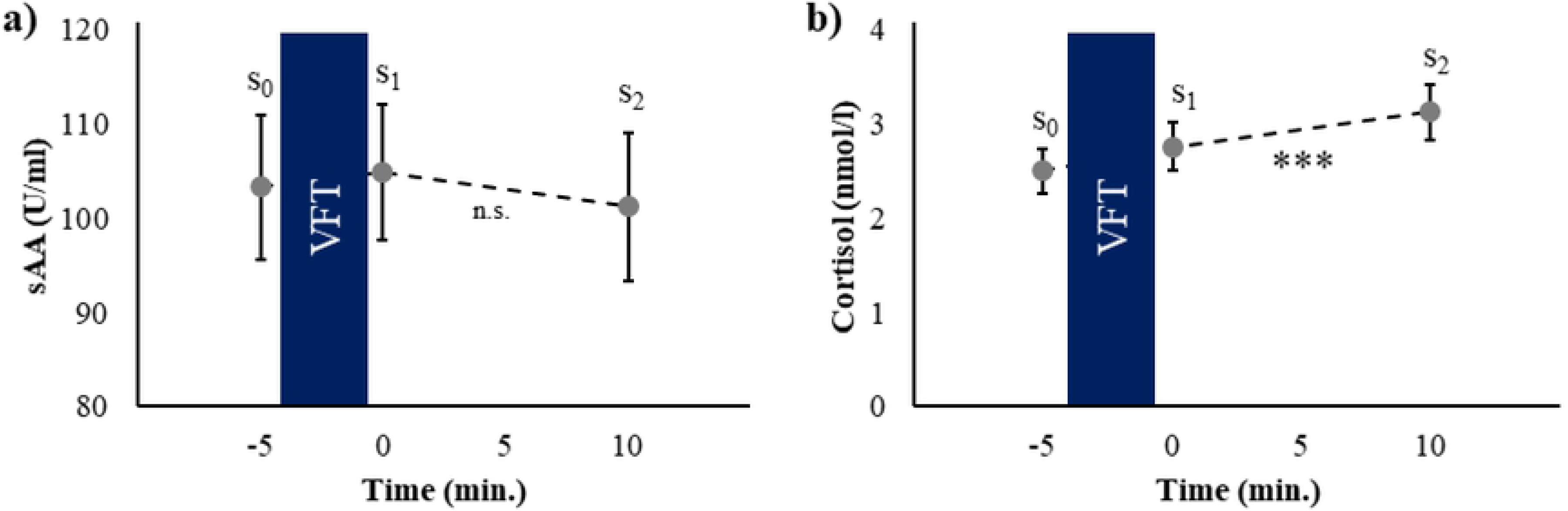
Alpha-amylase and cortisol responses. Time course of the salivary α-amylase (sAA, a) and cortisol (b) responses with respect to the verbal fluency task (VFT). Standard errors are shown as error bars.

### Cortisol

Cortisol levels significantly increased between t_0_ and t_1_ and increased further between t_1_ and t_2_ (*F*_(2, 168)_ = 19.29, *p* < .001, η_p_^2^ = .19, Fig 2b). However, cortisol levels were not associated with VFT performance (all *p* ≥ .370).

### Blood pressure and heart rate

When all participants were included into the analysis, a significant decrease in heart rate between t_0_ and t_2_ was found (*F*_(2, 159.8)_ = 6.36, *p* = .002, η_p_^2^ = .06, Fig 3a). Heart rate was not related with VFT performance (all *p* > .288). When only the non-hypertonic participants were included into the analyses, no effects were found (all *p* ≥ .069).

For the non-hypertonic participants, a main effect of the factor time was found for blood pressure (*F*_(1.87, 106.82)_ = 3.27, *p* = .041, η_p_^2^ = .05), indicating a decrease in blood pressure. Furthermore a main effect of the factor component was found (*F*_(1, 57)_ = 1610.19, *p* < .001, η_p_^2^ = .97), reflecting higher values for the systolic than the diastolic blood pressure component. No interaction between both factors (time x component) was found (*p* = .755, Fig 3b). Neither systolic nor diastolic blood pressure were related with VFT performance (all *p* ≥ .534). The same effects were found when including all participants into the analyses (time: *F*_(1.84, 150.65)_ = 5.59, *p* = .006, η_p_^2^ = .06; component: *F*_(1, 82)_ = 1549.48, *p* < .001, η_p_^2^ = .95; time x component: *p* = .086; correlations with VFT_perf_: all *p* ≥ .109).

**Fig 3.**
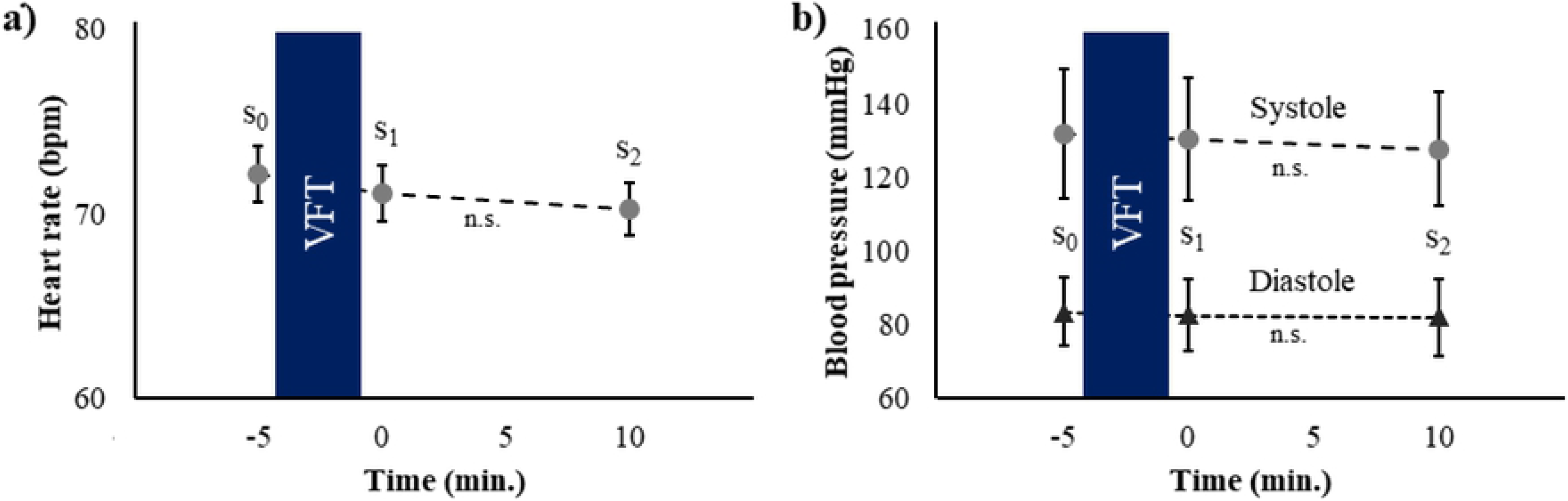
Heart rate and blood pressure responses. Time course of heart rate (in beats per minute (bpm), a) and blood pressure (b) with respect to the verbal fluency task (VFT). Standard errors are shown as error bars for heart rate and standard deviations for blood pressure.

### Subjective stress and affect

For the subjective ratings, a main effect of time (*F*_(2, 168)_ = 42.69,p < .001, η_p_^2^ = .34), a main effect of state (*F*_(1.45, 121.75)_ = 6.64, *p* = .005, η_p_^2^ = .07) and an interaction time x state (*F*_(3.0, 251.68)_ = 40.08, *p* < .001, η_p_^2^ = .32; Fig. 4a) were found. Post-hoc rmANOVAs showed that all states significantly changed during the experiment, i.e. that a main effect of time was found (stress: *F*_(2, 168)_ = 56.00, *p* < .001, η_p_^2^ = .40, effort: *F*_(2, 168)_ = 7.17, *p* = .001, η_p_^2^ = .08, tiredness: *F*_(2, 168)_ = 48.50, *p* < .001, η_p_^2^ = .37). Post-hoc t-tests showed that subjective stress perception significantly increased between t_0_ and t_1_ (*t*_(84)_ = −8.61, *p* < .001, *d* = 1.00, M_t0_ = 3.05 ± 1.75, M_t1_ = 4.73 ± 2.0) and decreased between t_1_ and t_2_ (*t*_(84)_ = 9.80, *p* < .001, *d* = 1.01, M_t1_ = 4.73 ± 2.00, M_t2_ = 2.95 ± 1.73). Subjective effort significantly increased between t_0_ and t_1_ (*t*_(84)_ = 4.30, *p* < .001, *d* = 0.46, M_t0_ = 3.66 ± 2.00, M_t1_ = 3.09 ± 1.85). Tiredness significantly increased between t_0_ and t_1_ (*t*_(84)_ = −9.60, *p* < .001, *d* = 1.32, M_t0_ = 2.22 ± 1.45, M_t1_ = 3.93 ± 2.03) and decreased between t_1_ and t_2_ (*t*_(84)_ = 6.40, *p* < .001, *d* = 0.82, M_t1_ = 3.93 ± 2.03, M_t2_ = 2.61 ± 1.57). Tiredness at t_1_ was negatively correlated with task performance (*r*_(84)_ = −.33, *p* = .002; Fig. 4b). None of the other subjective ratings was correlated with VFT performance after correction for multiple comparisons (all *p* ≥ .044).

Immediately before t_2_, the PANAS was filled-out by the participants. At this time point, positive affect was significantly higher than negative affect (*t*(77) = 16.35, *p* < .001, *d* = 1.5, M_positive_ = 3.03 ± 0.71, M_negative_ = 1.38 ± 0.41). After correction for multiple comparisons, a marginally significant negative correlation between negative affect and VFT performance was found (*r*_(77)_ = −.27, *p* = .017). Neither positive nor negative affect was related with any of the other physiological or psychological variables at t_2_ (all *p* ≥ .053).

**Figure 4.**
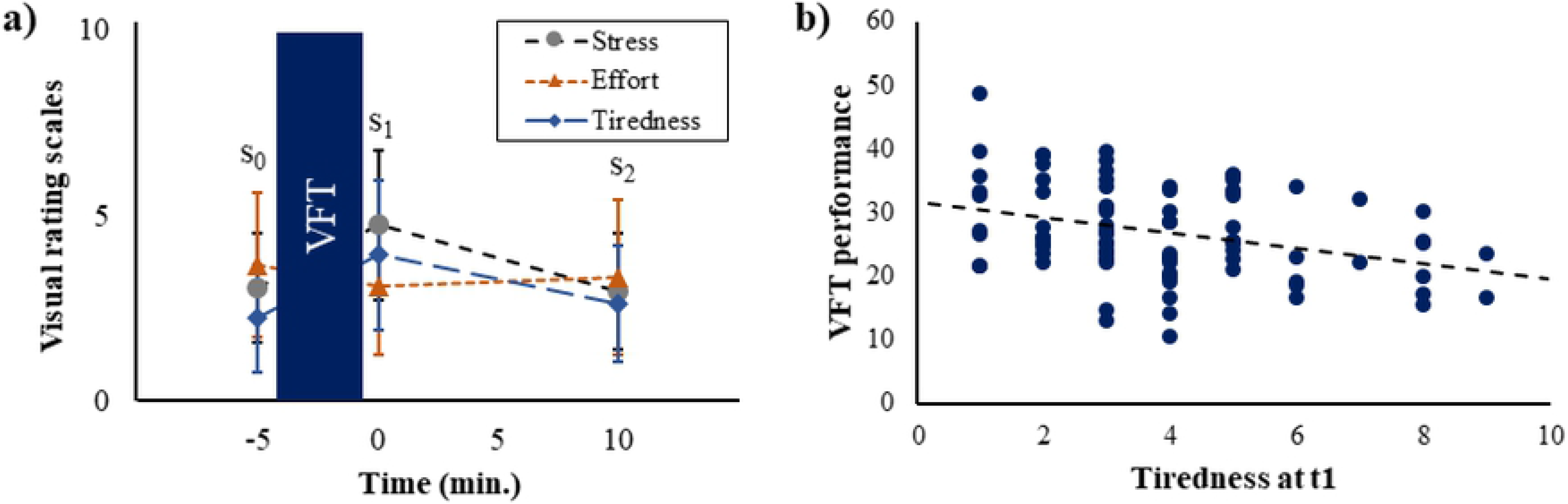
Subjective ratings and significant association with performance. Time course of subjective ratings for stress, effort, and tiredness (a) with respect to the verbal fluency task (VFT, a). In b), the association between VFT performance and tiredness at t_1_ is shown. Standard deviations are shown as error bars in a).

### Exploratory analysis

#### Depression

Since it has been shown previously that the stress response and cognitive performance can be associated with depression (e.g., 29), it was investigated whether this can be found in our study as well. Correlation analyses indicated that neither the physiological variables nor the cognitive performance were associated with depression (all *p* ≥ .113). Only negative affect and level of effort at t_0_ were related with depression (negative affect: *r*_(77)_ = .45, *p* < .001, effort at t_0_: *r*_(79)_ = .38, *p* = .001). After classifying the participants into depressed and non-depressed according to the cut-off value by Stein and Luppa (2012; 22), only a difference in negative affect was found between the groups (*t*(75) = −4.25, *p* < .001, *d* = 1.4, M_non-depressed_ = 1.3 ± 0.32, M_depressed_ = 1.9 ± 0.63). Neither the physiological variables nor the other subjective measures differed between depressed and non-depressed participants (all *p* ≥ .014).

#### Chronic stress

Next, it was investigated whether the stress response was associated with the level of perceived chronic stress during the last month. Subjective stress at t_0_ and t_1_ as well as negative affect were related with the PSS score (subjective stress at t_0_: *r*_(84)_ = .33, *p* = .002, subjective stress at t_1_: *r*_(84)_ = .34, *p* = .002, negative affect: *r*_(84)_ = .51, *p* < .001). Furthermore, a marginally significant association between VFT performance and PSS scores was found (*r*_(84)_ = .28, *p* = .008). None of the physiological variables was related to PSS levels (all *p* ≥ .105).

#### Time of day

Because a part of the experiment was conducted during a public event in the evening, and the rest was carried out in the morning or the afternoon, it was analyzed whether time of day was associated with task performance and with the stress response. All cortisol levels were associated with time of day (t_0_: *r*_(84)_ = −.54, *p* < .001, t_1_: *r*_(84)_ = −.50, *p* < .001, t_2_: *r*_(84)_ = −.42, *p* < .001, However, the percentage of the cortisol increase between t_0_ and t_2_ (i.e., the HPA axis response) was not related with time of day (*p* = .052). No associations were found for the other variables as well (all *p* > .003).

#### Anthropometric variables

At last, it was investigated whether the stress response in our experiment was associated with anthropometric factors (age, sex, and BMI) that are known to be typically related with the stress response. However, no associations were found when using the adjusted α-level (all *p* ≥ .003).

## Discussion

In our study, we have shown that a VFT can by itself induces an HPA axis response, i.e. an increase in cortisol levels. Furthermore, levels of subjective stress and effort as well as tiredness were higher immediately after the VFT than before. These effects (changes in HPA axis activity and subjective ratings) were independent of the cognitive performance during the task as well as of anthropometric and health variables. However, tiredness immediately after the task was negatively associated with task performance. Although cortisol levels were higher at later times of the day, no association between the HPA axis response and time of day was found.

Therefore, we suggest that a VFT can be an alternative to other stress induction methods such as the TSST (16,30) or the Montreal Imaging Stress Task (MIST; 31) which include combinations of social-evaluative and cognitive stressors, or the socially evaluated cold-pressor test (SECPT; 32–34) which involves a physiological and a social-evaluative stress component. In previous studies that used pure cognitive stressors, mathematical (e.g., 35,36) or inhibition tasks (e.g., the Stroop task; 37) have been used which also introduced physiological stress responses. We suggest that a VFT can be an easy to implement alternative to these cognitive stressors.

However, we cannot completely rule out that we created a social-evaluative component accidentally. This could have happened through the dictaphone that was placed in front of the participants or through the presence of our assistants.

Associations between HPA axis activity and VF performance have been reported previously. Greendale and colleagues (2000; 38) found that basal cortisol levels predicted VF performance four years later, i.e. higher cortisol levels were associated with worse verbal fluency performance. Fiocco and colleagues (2007; 39) reported that participants with lower educational levels showed higher cortisol responses to a TSST and performed worse in a VFT than participants with higher educational levels. Matsuo and colleagues (2003; 40) found reduced prefrontal activity during a VFT in participants with posttraumatic stress disorder (PTSD). However, they did not find performance differences between the PTSD patients and a healthy control group.

Our study is subject to some limitations that should be addressed in future research. First, other physiological variables (e.g., heart rate variability and electrodermal activity) should be measured. Furthermore, brain activity (e.g., assessed by means of EEG recordings) and its association with the stress response should be investigated. Second, longer time intervals should be used in future research to assess the time point and peak amplitude of the maximal cortisol increase as well as HPA axis recovery. Third, other control variables (e.g., childhood traumata or general cognitive performance) should be included in future studies. Last, associations between the HPA axis response and variations of the VFT (e.g., using a letter fluency task or using more categories) should be examined further.

## Conclusions

We conclude that a VFT is an acute stressor that induces a cortisol response without the need of further (e.g., social-evaluative) stress components. Therefore, it is a less time-consuming alternative to other stress tasks that can be used in field studies without much effort.

## Acknowledgement

We thank the visitors of the ‘Lange Nacht der Wissenschaften 2019’ for their interest in our work and for participating in this study. Furthermore, we thank Isabella Critelli, Aylin Gögsen, Emely Reyentanz, Luisa Schramm, and Theresa Walter for data collection.

